# Neural network-based alterations during repeated heat pain stimulation in major depression

**DOI:** 10.1101/553842

**Authors:** Edda Bilek, Zhenxiang Zang, Isabella Wolf, Florian Henrich, Carolin Mößnang, Urs Braun, Rolf-Detlef Treede, Walter Magerl, Andreas Meyer-Lindenberg, Heike Tost

## Abstract

The current study aimed to identify alterations in brain activation and connectivity related to nociceptive processing and pain sensitization in major depressive disorder (MDD), using repeated heat pain stimulation during functional magnetic resonance imaging (fMRI) in 37 MDD patients and 33 healthy controls. Regional activation did not differ between groups, but functional connectivity was significantly decreased in MDD in a neural network connecting frontal, temporal and occipital areas (family-wise error (FWE)-corrected pFWE = 0.045). Supplemental analyses suggested a significant association between network connectivity and trait neuroticism (p = 0.007) but not with the clinical state or familiar risk of MDD (all p values > 0.13). Our data relate a network-based phenotype for altered pain processing and antinociceptive control to MDD and encourage future studies on the shared intermediate neural psychological risk architecture of MDD and chronic pain.

## 1. Introduction

Depression and chronic pain are highly comorbid, as evidenced by pain reports in 66% of MDD patients (Arnow et al., 2006) and a two- to fourfold increase in depression rates in chronic pain (Gureje et al., 2008). These data suggests that both conditions may share common neurobiological mechanisms, but their nature and origins are poorly understood.

Patients with MDD frequently report more pain than controls, but are usually less sensitive to exteroceptive pain stimuli across modalities at threshold and tolerance limit (Bar et al., 2011; Dickens et al., 2003). Brain functional alterations during nociceptive processing and pain sensitization, which are well established for chronic pain (Apkarian et al., 2005), could underlie these abnormalities. However, the question of whether such neural deficits are present in MDD is underresearched. Few studies have specifically tackled this question using repeated heat pain stimulation during fMRI (Rodriguez-Raecke et al., 2014; Strigo et al., 2008). The authors linked decreased activations in right operculum and anterior cingulate cortex (ACC) during nociceptive processing and decreased brain stem activation during within-session pain sensitization to MDD, consistent with a disturbed processing of the emotional aspects of pain and a failure of the brain antinociceptive system (Rodriguez-Raecke et al., 2014).

For the origins of the observed neural system-level deficits in nociceptive processing, several factors appear plausible. First, the current clinical state or severity of MDD may play a role, calling for a comparison of the neuroimaging readouts between acute and remitted patients. Second, the deficits may relate to the genetic liability to depression, which can be explored by comparing healthy first-degree relatives (carrying genetic risk variants for MDD) to controls with a negative psychiatric family history. Third, a wealth of literature suggests a considerable overlap in personality traits conveying risk for both MDD and chronic pain, in particular qualities relating to trait neuroticism (e.g., trait anxiety, pain catastrophizing) (Gracely et al., 2004; Kadimpati et al., 2015). However, the effects of clinical state and risk factors on neural sensory pain processing in MDD are unexplored to date.

We used a well-established repeated heat pain protocol during fMRI to probe the questions of whether ***a)*** activation differences in key neural areas related to pain processing and short-term pain plasticity are present in MDD and ***b)*** whether these alterations involve network-based functional modifications in the brain nociceptive connectome. In supplementary analyses we further aimed to explore whether the uncovered alterations are influenced by ***c)*** the clinical state of the disorder, ***d)*** the familial liability to MDD, and/or ***e)*** trait neuroticism.

## 2. Experimental procedures

### 2.1. Participants

We recruited 70 individuals including 37 patients with MDD (mean age ± SD: 34.27 ± 11.19 years, 17 males) and 33 healthy controls without a history of mental illness (25.97 ± 9.86 years, 17 males). Psychiatric diagnoses were confirmed by trained clinical interviewers using the SCID-IV interview (First et al., 2001). From the MDD patients, 23 subjects were remitted while 14 individuals were in an acute disease phase. From the controls, 21 individuals had a negative psychiatric family history while 12 subjects were healthy first-degree relatives of MDD patients. Beck Depression Inventory (BDI) scores differed significantly between MDD and controls (15.27 ± 13.74 vs. 2.45 ± 3.85, *p* < 0.001) and the remitted and acute MDD patients (8.11 ± 8.01 vs. 22.05 ± 14.13, *p* < 0.002). Exclusion criteria included MRI contraindications, a history of neurological illness and current alcohol or drug abuse. All participants provided written informed consent for a protocol approved by the Ethics Committee of the University of Heidelberg.

### 2.2. Repetitive heat pain stimulation during fMRI

Brain nociceptive network challenge was achieved by repetitive heat pain stimulation during fMRI delivered with a 27 mm diameter combined heat foil Peltier thermode attached to the left volar forearm (Pathway, MEDOC, Ramat Yishai, Israel). The protocol was specifically designed for neuroimaging with a symmetrical on-off-pattern of stimuli: 60 stimuli of 48°C (ON) with 5s plateau with steep rises and declines at 8°C/s alternating with 5s of 32°C (OFF) (Jurgens et al., 2014). Total task duration was 17.6 min or 590 whole-brain scans. For fMRI analysis, the stimulus series was subdivided into 5 blocks of 12 ON and 12 OFF periods. For analysis of pain time course, a 0-100 numerical rating scale (NRS) was used; to avoid movement artefacts, ratings were obtained retrospectively for 10 evenly spaced time windows. Hyperalgesia was assessed 45-60min post-scan using 3 stimuli short increase to 50°C (no plateau) with immediate descent each to stimulation and control area (Peltier thermode, Medoc TSA II).

### 2.3. fMRI data acquisition and processing

See supplemental information for details.

### 2.4. Activation analysis

Activation analysis consisted of a two-level procedure. At the first level, we defined a general linear model for each subject including boxcar reference vectors for the two task conditions (convolved with the standard hemodynamic response function) and the head motion parameters. We high-pass filtered the data (cut-off: 128 seconds) and calculated individual contrast maps for pain processing (“pain>control”) and hyperalgesia development (parametric modulation of “pain>control” with time). Second-level statistical inference included univariate ANCOVA models with group (MDD vs. controls) as a factor, covarying for age and sex. Following prior work (Rodriguez-Raecke et al., 2014), statistical significance was assessed at *p* < 0.05, FWE corrected for multiple comparisons in an *a priori* defined mask covering bilateral ACC, insula, brain stem, thalamus, operculum and DLPFC (the latter defined as Brodman areas 9 and 46) using the the Automated Anatomical Labeling atlas (Tzourio-Mazoyer et al., 2002).

### 2.5. Construction of connectivity matrices and network-based statistic (NBS)

The functional network analysis followed established procedures (Cao et al., 2016; Cao et al., 2014; Zang et al., 2018). First, we parcellated the brain in 270 functional sphere ROIs of 5 mm diameter each. The node definitions included the 264 locations of the functional brain atlas by Power and colleagues (Power et al., 2011), to which we added 6 nodes located in bilateral caudate, amygdala and hippocampus (Cao et al., 2014). We extracted the mean node time series, regressed out white matter, cerebrospinal fluid and head movement and filtered the data with a 0.008 Hz high-pass filter. We then computed pairwise Pearson correlation coefficients between the processed node time series, which resulted in 270 × 270 matrices for each subject. Suprathreshold links were identified with an ANCOVA model (factor: MDD vs. controls, covariates: age and sex, initial threshold: *p* ≤ 0.0005, uncorrected) and we used NBS to identify clusters of node links that significantly differed between groups (Zalesky et al., 2010). The corrected cluster significance was defined by 5000 permutations.

### 2.6. Supplemental analyses

We conducted four supplemental analyses to probe the basis of the identified network hypoconnectivity during pain processing (see results section). To increase sensitivity, we performed post-hoc statistical analyses of the link estimates of the identified NBS network in SPSS (IBM SPSS Statistics for Windows, v25.0). To explore the brain network for ***a)*** potential group differences related to the development of hyperalgesia, we extracted the mean connectivity from the identified NBS network links for each stimulation block and participant and used a repeated measures ANOVA model with group (MDD vs. controls) and time (heat stimulation block 1-5) as factors to test for interaction effects. To further explore the brain network for potential group differences related to ***b)*** disease state (acute MDD vs. remitted MDD) and ***c)*** familial risk for MDD (healthy first-grade relatives vs. controls with a negative family history) and ***d)*** associations to trait neuroticism (STAI-T, (Laux et al., 1981)), we subjected the mean connectivity estimates of the identified NBS network links to independent t-tests (*b+c*) and correlation analysis (*d*).

## 3. Results

### 3.1. Heat-evoked pain

Repetitive heat stimulation elicited comparable levels of heat pain in healthy controls and MDD patients (45.5 ± 4.6 vs. 47.9 ± 3.3 NRS, *p* = 0.30) gradually rising during repeated stimulation (one-way ANOVA: *F*_2,58_ = 2.38, *p* < 0.05, ***Figure 1a***). Although group by time course interaction analysis was not significant, heat pain ratings increased significantly from the first time window in MDD patients (*p* < 0.005) reaching a peak and gradually declining thereafter (cubic trend: F_1,35_ = 6.31, *p* < 0.017), but not in healthy controls (all *p* values > 0.16; cubic trend: *F*_1,35_ = 2.60, *p* < 0.117). Testing heat pain 45-60 min later outside of the MRI scanner revealed significant heat hyperalgesia of the conditioned skin in both groups (both p<0.001, ***Figure 1b***).

**Legend Figure 1.**
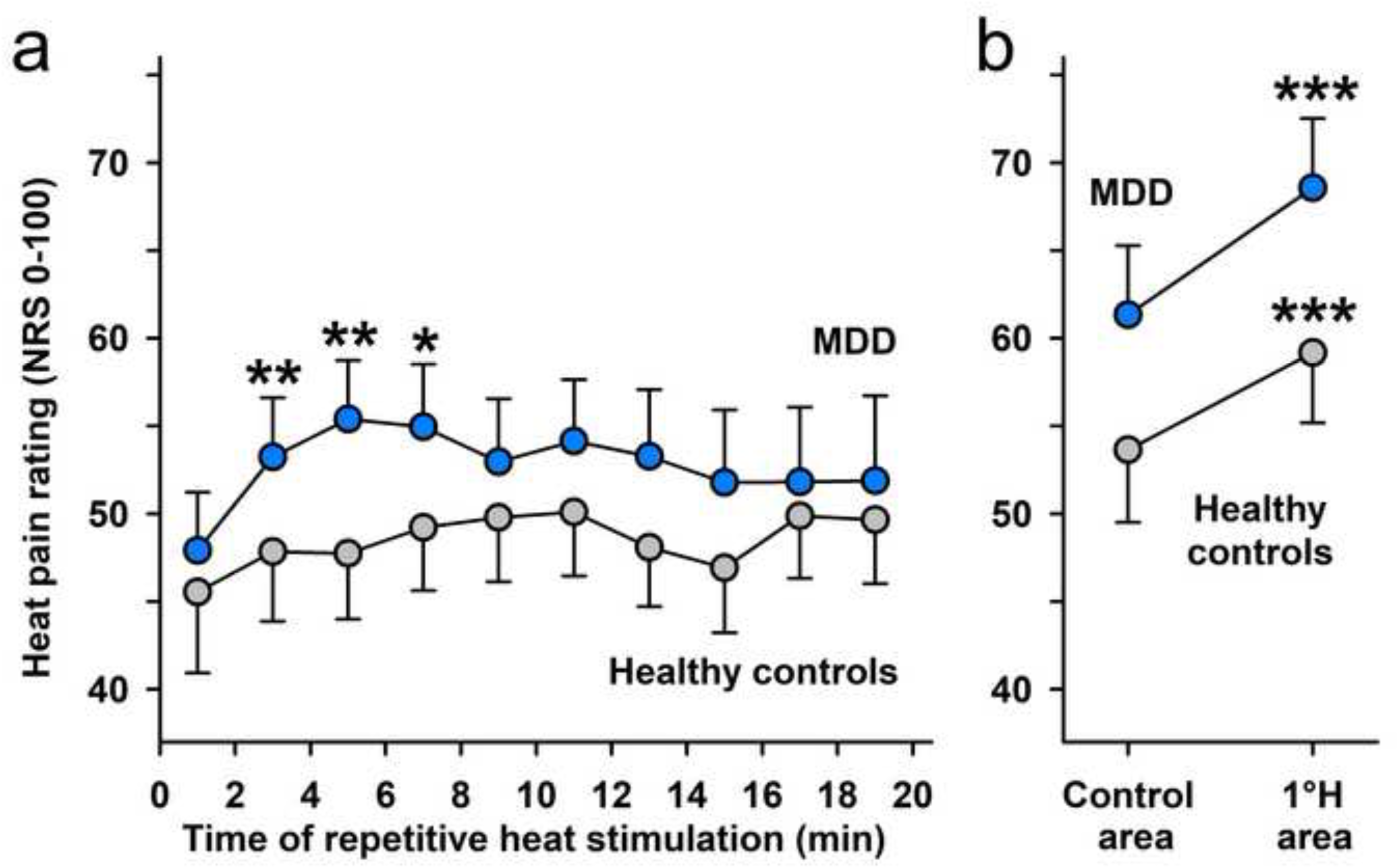
***a)*** Repetitive noxious heat stimulation elicits moderate heat pain with slowly increasing magnitude over time. MDD patients exhibit a more pronounced increase of heat pain over time (overshoot of heat hyperalgesia in early time windows) compared to healthy controls; * p<0.05 and ** p<0.01 vs. first pain rating interval. ***b)*** Repetitive heat stimulation elicits heat hyperalgesia in the conditioned skin area at 45-60 min after conditioning (primary hyperalgesia 1°H); *** p<0.001 vs. contralateral unconditioned (control area on right volar forearm). MDD, major depressive disorder; NRS, numerical rating scale.

### 3.2. Activation analysis

We detected a highly significant activation increase in brain regions associated with pain processing including bilateral insula, aMCC, supplementary motor area and thalamus as well as a highly significant activation increase over time in key control areas of the nociceptive network including bilateral DLPFC (all *p* values < 0.05, FWE-corrected for the whole brain, see ***Figure 2a and b*** and ***Supplementary Tables S1 and S2*** for a full list of significant regions). However, we detected no significant activation differences between MDD patients and controls for pain processing or the development of hyperalgesia in the *a priori* defined regions (all *p* values *>* 0.05, FWE corrected for region of interest). Moreover, subsequent data inspection at exploratory statistical thresholds (*p* < 0.001, k = 50, uncorrected for multiple comparisons) did not provide evidence for a sizeable group difference in any other brain region.

**Legend Figure 2.**
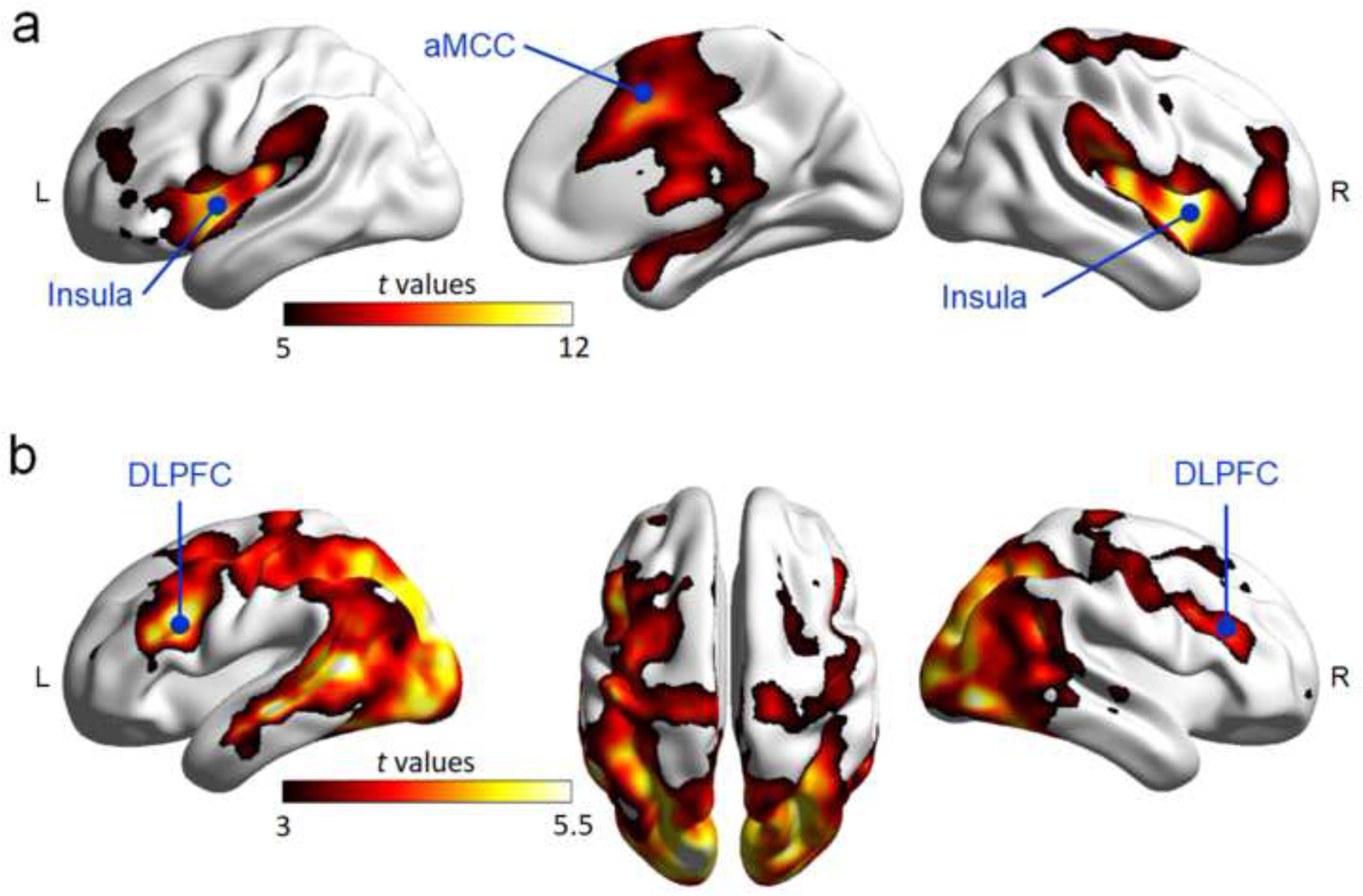
***a)*** Repetitive heat pain stimulation during fMRI (contrast: pain>control) elicits highly significant activations (*p*_FWE_ < 0.05, whole-brain corrected) in key cortical areas of nociceptive processing including bilateral insula and anterior midcingulate cortex (aMCC). ***b)*** Over time (contrast: [pain>control]*time, parametric modulation analysis), heat pain stimulation results in a highly significant increase of activation (*p*_FWE_ < 0.05, whole-brain corrected) in key control areas of nociceptive processing including bilateral dorsolateral prefrontal cortex (DLPFC). For illustration purposes, we displayed activation maps at a threshold of *t* = 3 using BrainNet Viewer (https://www.nitrc.org/projects/bnv/). fMRI, functional magnetic resonance imaging; FWE, family wise error.

### 3.3. Network-based statistic (NBS)

NBS identified a cluster of links with a significant decrease in functional connectivity in MDD compared to controls during pain processing (cluster-level FWE-corrected *p* = 0.045, ***Figure 3a and b***). The cluster consisted of 65 nodes and 70 links mainly interconnecting frontal, temporal and occipital cortices. Specifically, the majority of links (53%) included at least one node interconnecting the frontal cortex with other areas. A detailed description of all network nodes and links of the cluster is provided in ***Supplementary Table S3***.

**Legend Figure 3.**
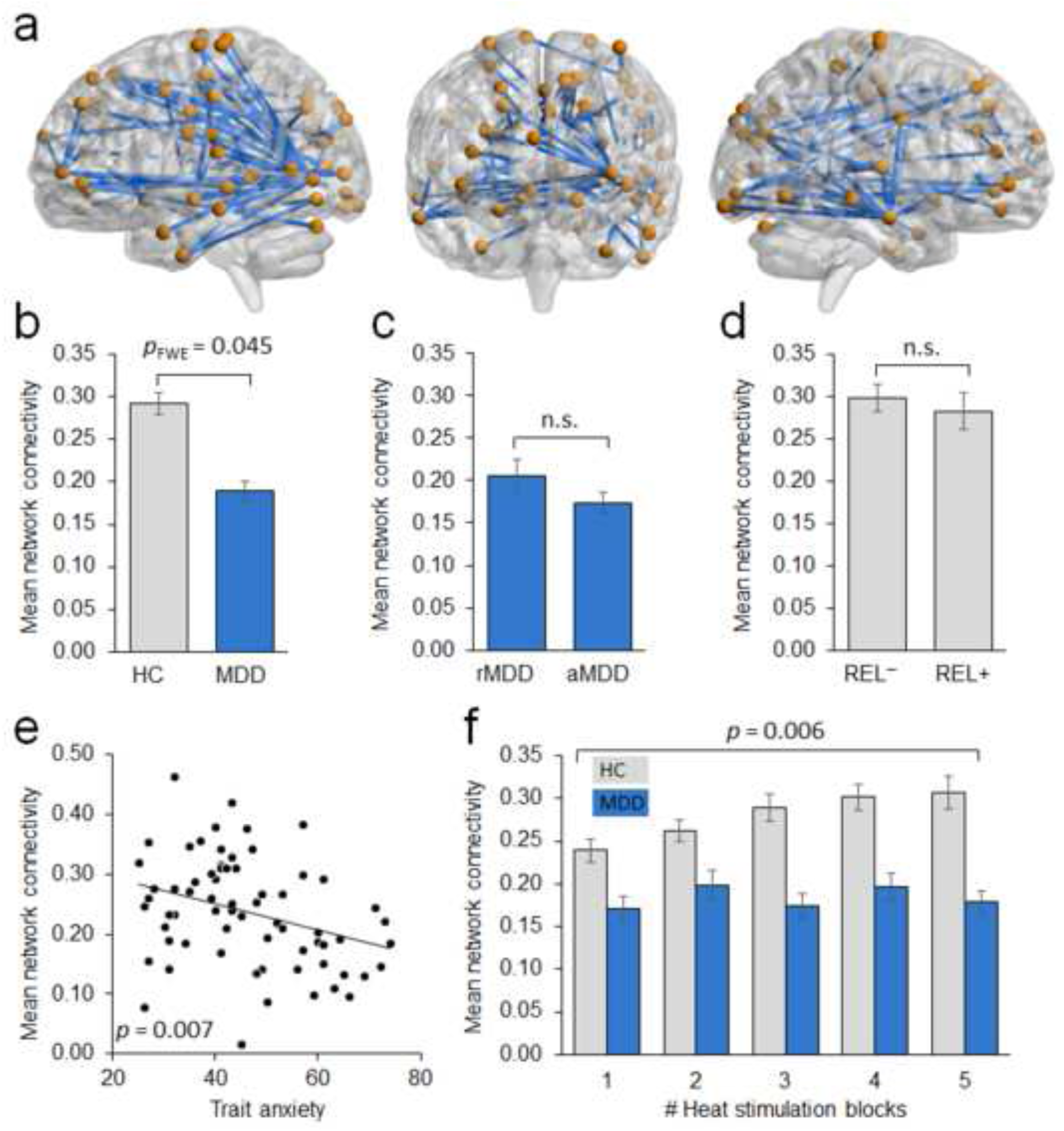
***a)*** Cortical cluster of identified node links superimposed on a 3D brain template ***b)*** demonstrating a significant connectivity decrease in MDD patients compared to healthy controls during fMRI repetitive heat pain stimulation (*p*_FWE_ < 0.045, cluster permutation corrected). ***c)*** No evidence for differences in cluster connectivity in acute vs. remitted MDD patients or ***d)*** healthy controls with vs. without a first-degree MDD relative but ***e)*** significant correlation of connectivity estimates and trait anxiety scores of participants (*p* = 0.007). ***f)*** Significant group by time interaction effect in cluster connectivity in healthy controls vs. MDD patients over time (*p* = 0.006). HC, healthy control subject; MDD, major depressive disorder; rMDD, remitted MDD; aMDD, acute MDD; REL-, no family history of MDD; REL+, healthy MDD relative; fMRI, functional magnetic resonance imaging; FWE, family wise error.

### 3.4. Supplemental analyses

In post-hoc statistical analyses of the identified NBS network we detected no group differences in network connectivity in acute vs. remitted MDD patients (*t* = 1.52, *p* = 0.13, ***Figure 3c***) or healthy controls without a psychiatric family history vs. those with a first-degree relative with MDD (*t* = 0.10, *p* = 0.92, ***Figure 3d***). However, we observed a significant negative correlation between the functional connectivity estimates of the identified network and the trait anxiety of participants (*r* = −0.32, *p* = 0.007, ***Figure 3e***). In addition, we detected a significant group by time interaction effect with a significant increase in the functional connectivity of the identified NBS network in healthy controls and a relative absence of this increase in MDD (*F*_(4,272)_ = 3.73, *p* = 0.006, ***Figure 3f***).

## 4. Discussion

In this study, we used a well-established repeated heat pain protocol during fMRI to study nociceptive processing and pain sensitization in depression and explore the potential origins of the identified neural alterations (Jurgens et al., 2014). Our findings did not provide evidence for regional brain activation deficits in MDD during pain processing. Instead, we identified connectivity alterations in a neural network of frontal, temporal and occipital nodes displaying deficient connectivity. Supplemental analyses suggested that the network deficits related to trait anxiety, but not the clinical severity of MDD or the genetic liability to the disorder in unaffected individuals.

In contrast to a prior study with a similar protocol (Rodriguez-Raecke et al., 2014), we did not detect any significant brain activation differences related to nociceptive processing or within-session pain sensitization in MDD, not even at an uncorrected exploratory significance level. We may speculate that differences in the experimental protocols explain this discrepancy. While Rodriguez-Raecke and coworkers acquired pain intensity ratings during fMRI, we refrained from such assessments to minimize higher-order cognitive processes related to the evaluation of pain during fMRI. Thus, while their conclusions on disturbed emotional processing of pain and failure of the antinociceptive system in MDD may still be valid, the corresponding brain activation findings may be induced by pain protocols engaging higher-order cognitive processes such as pain appraisal.

We further used NBS to identify a cortical network with reduced functional connectivity of node links during pain processing in MDD. We have repeatedly demonstrated the sensitivity and robustness of this method for the identification of neural connectivity risk markers for psychiatric disorders (Cao et al., 2016; Zang et al., 2018). Notably, although widespread in nature, the majority of links of the identified cluster involved node definitions mapping to the frontal cortex, which points to a potential regulatory role of the functional network during pain processing. Previous investigation has shown altered prefrontal-limbic activation patterns in MDD patients to pain anticipation (Strigo et al., 2008). Consistent with this, the detected network hypoconnectivity profile in MDD related to the absence of the functional connectivity increase with continuing pain stimulation displayed by the healthy controls. This is consistent with the electroencephalography finding of a lower increase in network redundancy parameters in MDD patients after electrical pain stimulation (Leistritz et al., 2013). These data suggest that the identified cortical network reflects increased coupling of the brain antinociceptive system during pain sensitization, which appears to be absent in MDD, but exaggerated in borderline personality disorder (Schmahl et al., 2006).

Our supplemental analyses on the potential origins of the identified neural system-level deficit revealed a significant correlation between the functional connectivity estimates of the assumed nociceptive control network and trait anxiety, an important dispositional risk factor for MDD related to dysfunctional prefrontal-limbic regulation of affective responses (Pezawas et al., 2005). Trait anxiety shows significant conceptual overlap with pain catastrophizing, a well-established maladaptive cognitive style that is related to negative affectivity in trait-anxious and depressed patients and a potent psychological risk factor for the development of chronic pain (Malfliet et al., 2017; Thieme et al., 2015; Treede et al., 2018).

Taken together, our findings identify a network-based functional phenotype for altered pain processing and antinociceptive control in MDD that is unrelated to the clinical state and familial disposition for the disorder. Instead, we provide evidence for the role of an intermediate psychological risk factor for both MDD and chronic pain related to negative affectivity and linked to a higher-order cortical control network involving the prefrontal cortex. These data call for future research examining neural network-based functional alterations in prefrontal regulatory circuits in MDD (Diener et al., 2012) and encourage the study of the shared temperamental risk architecture of depression and chronic pain.

## Contributors

WM, HT, RDT and AML designed the study. IW, FH and EB collected the data. EB, ZZ, FH, WM, CM, and UB conducted the analyses. EB, ZZ and HT wrote the first draft of the manuscript. All authors contributed to and have approved the final manuscript.

## Acknowledgements

The research leading to these results was supported by the German Research Foundation within the Collaborative Research Center (SFB) 1158 (project B09 and S01). The authors thank Axel Schäfer, Ilka Alexi, Roland Neubauer, Michelle Berkemann, Marlena Itz and Yuri Sevchenko for valuable research assistance.

## References

Apkarian, A.V., Bushnell, M.C., Treede, R.D., Zubieta, J.K., 2005. Human brain mechanisms of pain perception and regulation in health and disease. Eur J Pain 9, 463–484.

Arnow, B.A., Hunkeler, E.M., Blasey, C.M., Lee, J., Constantino, M.J., Fireman, B., Kraemer, H.C., Dea, R., Robinson, R., Hayward, C., 2006. Comorbid depression, chronic pain, and disability in primary care. Psychosom Med 68, 262–268.

Bar, K.J., Terhaar, J., Boettger, M.K., Boettger, S., Berger, S., Weiss, T., 2011. Pseudohypoalgesia on the skin: a novel view on the paradox of pain perception in depression. J Clin Psychopharmacol 31, 103–107.

Cao, H., Bertolino, A., Walter, H., Schneider, M., Schafer, A., Taurisano, P., Blasi, G., Haddad, L., Grimm, O., Otto, K., Dixson, L., Erk, S., Mohnke, S., Heinz, A., Romanczuk-Seiferth, N., Muhleisen, T.W., Mattheisen, M., Witt, S.H., Cichon, S., Noethen, M., Rietschel, M., Tost, H., Meyer-Lindenberg, A., 2016. Altered Functional Subnetwork During Emotional Face Processing: A Potential Intermediate Phenotype for Schizophrenia. JAMA Psychiatry 73, 598–605.

Cao, H., Plichta, M.M., Schafer, A., Haddad, L., Grimm, O., Schneider, M., Esslinger, C., Kirsch, P., Meyer-Lindenberg, A., Tost, H., 2014. Test-retest reliability of fMRI-based graph theoretical properties during working memory, emotion processing, and resting state. Neuroimage 84, 888–900.

Dickens, C., McGowan, L., Dale, S., 2003. Impact of depression on experimental pain perception: a systematic review of the literature with meta-analysis. Psychosom Med 65, 369–375.

Diener, C., Kuehner, C., Brusniak, W., Ubl, B., Wessa, M., Flor, H., 2012. A meta-analysis of neurofunctional imaging studies of emotion and cognition in major depression. Neuroimage 61, 677–685.

First, M., Spitzer, R., Gibbon, M., Williams, J., 2001. Structured Clinical Interview for DSM-IV-TR Axis I Disorders, Research Version, Patient Edition with Psychotic Screen (SCID-I/P W/ PSY SCREEN). New York State Psychiatric Institute, New York.

Gracely, R.H., Geisser, M.E., Giesecke, T., Grant, M.A., Petzke, F., Williams, D.A., Clauw, D.J., 2004. Pain catastrophizing and neural responses to pain among persons with fibromyalgia. Brain 127, 835–843.

Gureje, O., Von Korff, M., Kola, L., Demyttenaere, K., He, Y., Posada-Villa, J., Lepine, J.P., Angermeyer, M.C., Levinson, D., de Girolamo, G., Iwata, N., Karam, A., Guimaraes Borges, G.L., de Graaf, R., Browne, M.O., Stein, D.J., Haro, J.M., Bromet, E.J., Kessler, R.C., Alonso, J., 2008. The relation between multiple pains and mental disorders: results from the World Mental Health Surveys. Pain 135, 82–91.

Jurgens, T.P., Sawatzki, A., Henrich, F., Magerl, W., May, A., 2014. An improved model of heat-induced hyperalgesia--repetitive phasic heat pain causing primary hyperalgesia to heat and secondary hyperalgesia to pinprick and light touch. PLoS One 9, e99507.

Kadimpati, S., Zale, E.L., Hooten, M.W., Ditre, J.W., Warner, D.O., 2015. Associations between Neuroticism and Depression in Relation to Catastrophizing and Pain-Related Anxiety in Chronic Pain Patients. PLoS One 10, e0126351.

Laux, L., Glanzmann, P., Schaffner, P., Spielberger, C.D., 1981. State-Trait-Angstinventar (STAI). Theoretische Grundlagen und Handanweisung. Beltz Test GmbH.

Leistritz, L., Weiss, T., Bar, K.J., De VicoFallani, F., Babiloni, F., Witte, H., Lehmann, T., 2013. Network redundancy analysis of effective brain networks: a comparison of healthy controls and patients with major depression. PLoS One 8, e60956.

Malfliet, A., Coppieters, I., Van Wilgen, P., Kregel, J., De Pauw, R., Dolphens, M., Ickmans, K., 2017. Brain changes associated with cognitive and emotional factors in chronic pain: A systematic review. Eur J Pain 21, 769–786.

Pezawas, L., Meyer-Lindenberg, A., Drabant, E.M., Verchinski, B.A., Munoz, K.E., Kolachana, B.S., Egan, M.F., Mattay, V.S., Hariri, A.R., Weinberger, D.R., 2005. 5-HTTLPR polymorphism impacts human cingulate-amygdala interactions: a genetic susceptibility mechanism for depression. Nat Neurosci 8, 828–834.

Power, J.D., Cohen, A.L., Nelson, S.M., Wig, G.S., Barnes, K.A., Church, J.A., Vogel, A.C., Laumann, T.O., Miezin, F.M., Schlaggar, B.L., Petersen, S.E., 2011. Functional network organization of the human brain. Neuron 72, 665–678.

Rodriguez-Raecke, R., Ihle, K., Ritter, C., Muhtz, C., Otte, C., May, A., 2014. Neuronal differences between chronic low back pain and depression regarding long-term habituation to pain. Eur J Pain 18, 701–711.

Schmahl, C., Bohus, M., Esposito, F., Treede, R.D., Di Salle, F., Greffrath, W., Ludaescher, P., Jochims, A., Lieb, K., Scheffler, K., Hennig, J., Seifritz, E., 2006. Neural correlates of antinociception in borderline personality disorder. Arch Gen Psychiatry 63, 659–667.

Strigo, I.A., Simmons, A.N., Matthews, S.C., Craig, A.D., Paulus, M.P., 2008. Association of major depressive disorder with altered functional brain response during anticipation and processing of heat pain. Arch Gen Psychiatry 65, 1275–1284.

Thieme, K., Turk, D.C., Gracely, R.H., Maixner, W., Flor, H., 2015. The relationship among psychological and psychophysiological characteristics of fibromyalgia patients. J Pain 16, 186–196.

Treede, R.D., Rief, W., Barke, A., Aziz, Q., Bennett, M.I., Benoliel, R., Cohen, M., Evers, S., Finnerup, N.B., First, M.B., Giamberardino, M.A., Kaasa, S., Korwisi, B., Kosek, E., Lavand’homme, P., Nicholas, M., Perrot, S., Scholz, J., Schug, S., Smith, B.H., Svensson, P., Vlaeyen, J.W., Wang, S.J., 2018. Chronic pain as a symptom or a disease: the IASP classification of chronic pain for the international classification of diseases ICD-11. accepted July 2018.

Tzourio-Mazoyer, N., Landeau, B., Papathanassiou, D., Crivello, F., Etard, O., Delcroix, N., Mazoyer, B., Joliot, M., 2002. Automated anatomical labeling of activations in SPM using a macroscopic anatomical parcellation of the MNI MRI single-subject brain. Neuroimage 15, 273–289.

Zalesky, A., Fornito, A., Bullmore, E.T., 2010. Network-based statistic: identifying differences in brain networks. Neuroimage 53, 1197–1207.

Zang, Z., Geiger, L.S., Braun, U., Cao, H., Zangl, M., Schafer, A., Moessnang, C., Ruf, M., Reis, J., Schweiger, J.I., Dixson, L., Moscicki, A., Schwarz, E., Meyer-Lindenberg, A., Tost, H., 2018. Resting-state brain network features associated with short-term skill learning ability in humans and the influence of N-methyl-d-aspartate receptor antagonism. Netw Neurosci 2, 464–480.

